# Integrating satellite data and field surveys to predict bird communities in the Peruvian Amazon

**DOI:** 10.1101/2025.07.17.665287

**Authors:** Andrew Christopher Slater, Chris A. Kirkby, Chris Ketola, Ian Hartley, Alex Bush

## Abstract

Tropical forests harbour some of the highest biodiversity on Earth but are undergoing rapid loss and degradation. In the Amazon more than one-third of forests have been altered through human activities, with major implications for wildlife communities. While Earth observation satellites effectively monitor forest cover at scale, it remains unclear how well satellite-derived variables capture variation in bird communities. We tested whether Landsat reflectance and vegetation indices can predict bird species occurrence and community composition in the Peruvian Amazon. We analysed 3,129 point counts and mist-net bird surveys conducted over 16 years in the Tambopata Forest, south-eastern Peru. As predictors we compared the effectiveness of remote sensing derived surface reflectance and vegetation indices (e.g. NDVI and tasselled cap), with traditional land-type and forest cover descriptors. Species occurrence probabilities and community composition of 135 frequently recorded bird species were estimated using multi-species occupancy models that account for imperfect detection. Models using Landsat reflectance and vegetation indices outperformed those based on habitat categories in predicting species occupancy (mean AUC = 0.68 vs 0.58). They also achieved high predictive accuracy (AUC > 0.7) for more species (49 compared with 20). However, low detection rates across surveys limited all models’ ability to accurately estimate full community composition and to detect change over time. Our results demonstrate that satellite-derived variables can improve predictions of bird occurrence compared with habitat categories, but their effectiveness depends strongly on survey design and species detectability. Integrating remote sensing with well-structured field surveys provides a scalable approach to monitoring biodiversity trends in tropical forests.

## 2 Introduction

Tropical forests are the most biodiverse areas on Earth (1,2), and support over a billion people with food, materials, and services (3). They also play a key role in the global carbon cycle, accounting for half of terrestrial uptake and two-thirds of forest carbon sinks in biomass, soil, and deadwood (4,5). However, clear-cut logging and commercial agriculture are altering these habitats, with over 17% of the Amazon Basin deforested and 38% of the remainder degraded (6–8). Conservation strategies, such as the Reducing Emissions from Deforestation and Degradation (REDD+) initiative, have generally reduced deforestation, enhanced carbon stocks, and improved sustainable forest management (9–11). However, half of the world’s tropical forest reserves continue to lose taxonomic and functional biodiversity due to disturbances beyond forest loss (12,13). While forest cover change can be mapped in near real-time (14), assessing the efficacy of conservation strategies requires evaluating the current condition of biodiversity within the ecosystem and where and how it changes over time (15–17).

Ecosystem structure is often described by categorical elements such as intact, logged, secondary, or plantation forest, yet these definitions can be ambiguous, subjective, and may not reflect the most effective descriptors of an ecosystem’s condition (18–20). In this regard, essential biodiversity variables, including species distribution, abundance, and community composition, have been proposed as additional criteria to monitor (21,22). Satellite remote sensing (RS) cannot directly see these variables, but it can provide broader, scalable environmental descriptors, alternative to field surveys, that act as proxies for essential biodiversity variables, such as ecological diversity, phenology and physiology (23–26). Continuous RS variables have been shown to outperform discrete land classes at describing plant species richness and diversity (27). Similar approaches have been applied to birds, where RS indices predict species richness and community composition (28,29), the structure of which has been used to assess the effectiveness of reforestation (30), and to show that they are negatively affected by patch size reduction within the Brazilian Amazon (31). RS surface reflectance values can therefore be used to infer vegetation and habitat structure as continuous gradients, avoiding the limitations of pre-defined classes. Yet, while RS variables can infer potential structure and ecological richness, there is little understanding of how they relate to the organisation of bird communities within tropical forests (32–34).

Biodiversity surveys provide essential information but are time consuming, expensive, and often limited in their spatial and temporal coverage (35,36). Linking survey data with RS descriptors in statistical models offers a way to estimate community and ecosystem structure across a landscape with limited expense (37). Because Amazonian bird communities are shaped by the characteristics of the forest in which they occur, we expect RS surface reflectance values to capture elements of these patterns. The continually updated nature of RS data also presents opportunities for this method to be used in monitoring ongoing changes in community structure. In this study, we test whether RS reflectance can better predict bird species occupancy than traditional habitat descriptors, and whether these models can predict observed community composition. We further hypothesise that survey replications and species detection rates will constrain model performance and the ability to detect temporal change. To evaluate these questions, we compare model predictive accuracy using species-level (AUC) and community-level (Bray-Curtis dissimilarity) metrics, on withheld data. We also assess how survey effort influences the power to monitor biodiversity trends. By bringing these elements together, we aim to provide a practical tool for policy makers and ecologists to assess the successes and failures of management strategies, and to support decisions about financial incentives.

## 3 Methods

### 3.1 Study area and surveys

Survey data were collected by FaunaForever (www.faunaforever.org) to monitor species diversity across landscapes in the Madre de Dios region, near the Tambopata River and Puerto Maldonado in south-eastern Peru (**Error! Reference source not found.**). All collection and subsequent use of these data complied with FaunaForever’s data use policies. Researchers and volunteers conducted bird surveys at 637 stations between 2004 and 2020. Bird surveys were conducted under permits issued by INRENA (Instituto Nacional de Recursos Naturales) and later SERNANP (Servicio Nacional de Áreas Naturales Protegidas por el Estado) for sites within the Tambopata National Reserve (RNTAMB; permits 20 S/C-2003-INRENA-IANP, 004-2010-SERNANP-RNTAMB-JEF, 014-2011-SERNANP-RNTAMB-JEF, 012-2012-SERNANP-RNTAMB-JEF). Surveys outside national protected areas were undertaken with the permission of landowners or concession title holders in the context of their approved management plans (DEMA). The stations were clustered around 25 centres that were located between 1.5km and 170km apart. Within centres, stations were between 200m and several kilometres apart and surveyed one to 35 times (median = 4), totalling 3,129 surveys. Stations surveyed in multiple years were treated as independent, since populations and forest structure can change over extended periods of time. This resulted in a total of 1,135 independent stations, which were surveyed between one and 21 times, with a mode of 1 survey, a median of 2 surveys and a mean of 2.9 surveys per site (SD ±2.4). Surveys used point counts (10-minute visual/audio records), or net counts (36m mist-net captures). Simulation studies have shown that models built using species with fewer than 10 observations are prone to unreliable predictions, but the precise minimum required varies with species’ range size and study context (38). Based on this evidence, we applied a pragmatic threshold, not intended as a universal standard, that balanced the inclusion of as many species as possible with the need for reliable model estimates. We therefore retained only species observed at more than 10 stations. Of the 358 species identified, this resulted in 138 species being used for model training.

**Fig 1:**
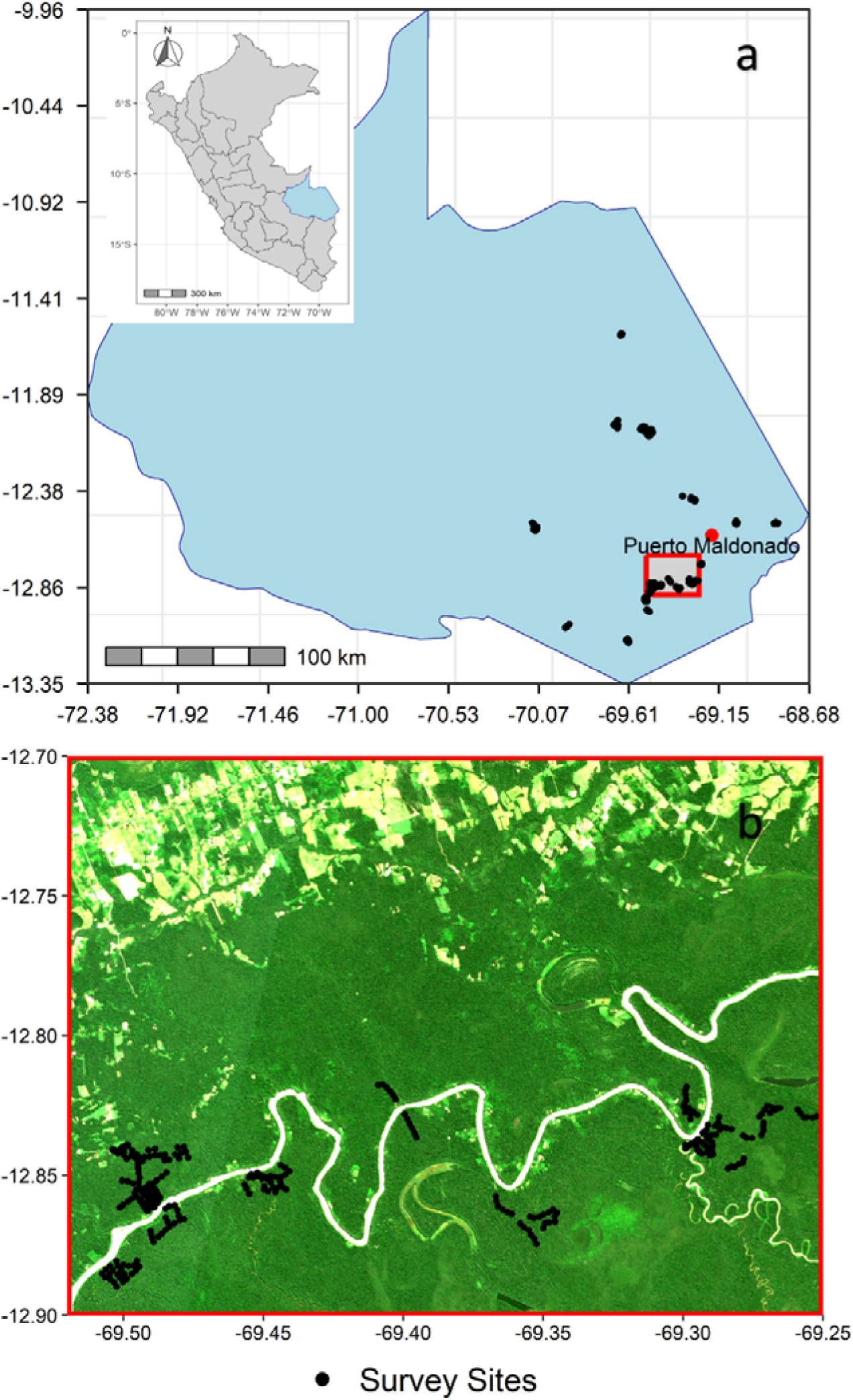
**(a)** The location of survey stations within the Madre de Dios region, south-eastern Peru. The inset map shows Peru with Madre de Dios highlighted in blue. The grey rectangle bounded in red indicates the area shown in panel **(b)**. Panel **(b)** illustrative zoom of the highlighted area showing local dispersion of survey stations. The satellite backdrop in (**b**) is provided for context only and was not used in analysis. Basemap data: Natural Earth (public domain). Panel (b) backdrop: Sentinel-2 L2A imagery (Copernicus, cc BY 4.0)(109).

### 3.2 Environmental variables

Habitat was classified by land class and forest degradation. Land classes (floodplain forest, terra firma forest, and agricultural) were based on the MapBiomas Peru Project, Collection**-1** of the Annual Land Cover and Land Use Series for Peru 2013; (39). MapBiomas classifications are based on Landsat imagery at 30m resolution using a random forest approach to produce annual land cover and land use maps across Peru. We selected the 2013 dataset as it represents the midpoint of our 2004-2020 survey period and provides a consistent baseline for all survey sites. While this inevitably simplifies temporal changes in land cover, our survey stations were largely in stable forested areas where major shifts were unlikely.

Forest degradation was defined using the Hansen et al. Global Forest Change dataset, which reports annual forest loss from 2000 onwards (14). Pixels with no reported loss were classified as primary forest, while those with any recorded loss were classified as secondary forest.. This binary classification is a coarse simplification of the successional gradient but was necessary to align with categories recorded in some field surveys, and to enable comparability across datasets. The proportion of non-primary forest, equating to forest assumed to have been deforested or degraded, was calculated within three different radii around each station (1km, 2km and 5km). No cross validation was performed between the MapBiomas and Hansen datasets, so forest classifications may not align. These limitations highlight the constraints of categorical land-cover datasets for ecological modelling, and partly motivate our parallel use of continuous remotely sensed reflectance variables to better capture environmental gradients.

Of the 1135 surveys, 489 were on terra firma, 603 on floodplain and 43 on agricultural land, and 1109 were defined as being inside primary forest, and 26 in secondary forest. Elevation can influence the distribution of tropical forest bird species (40,41). However, all survey points were within the narrow range between 175m and 310m elevation, and unlikely to affect model performance; it was therefore not included in the analysis.

### 3.3 Satellite derived variables

Reflectance signatures at different wavelengths can be employed to assess the greenness and moisture content of vegetation, with some correlation observed between these signatures and the leaf structure of a plant (42). Reflectance and water content descriptors help describe forest canopy physiology (43). No single metric captures canopy spectral diversity (44), and the spectral characteristics and spatial scales that best capture canopy variation are unknown, so various indices were used to describe it. While many satellite platforms now provide passive optical surface reflectance data, as well as active radar reflectance, the Landsat missions were the only platforms that covered the entire period of the field surveys (Landsat-5 for 2004-2011, Landsat-7 for 2012, and Landsat-8 for 2013-2020). For all missions we used USGS Collection-1 surface reflectance products, which provide consistent atmospherically corrected data across sensors. Frequent cloud cover over tropical forests can greatly reduce the number of available images that have sufficient clarity for analysis (45). To address poor visibility, annual cloud-masked, median pixel value images were created in Google Earth Engine. Annual composites necessarily average across all available cloud-free imagery within each year and therefore do not represent the specific timing of individual bird surveys. While this approach may mask seasonal changes such as leaf flushing and fruiting cycles, or logging and burning, phenological variation in tropical forests is generally lower than in temperate ecosystems (26,46,47). Each image contained six reflectance bands, which were used to calculate seven vegetation indices: the normalised difference vegetation index (NDVI), enhanced vegetation index (EVI), normalised burn ratio (NBR), normalised difference water index (NDWI), and the trio of tasselled cap wetness, greenness, and brightness (TCW, TCG, TCB). Both NDVI and EVI were used to detect vegetation cover and vigour. NDVI was included as one of the most commonly used indices, but it is prone to saturation in dense forests such as the Amazon, whereas EVI is better able to penetrate the canopy and capture sub-layers. We used NBR to highlight freshly opened soil, and NDWI to highlight water content within the canopy. Tasselled cap indices provide an alternative way of describing reflectance and structure, and have been shown to correlate with, but show more detail than, NDVI (48).

For each annual survey, the mean and standard deviation of pixels in each of the reflectance bands and indices were calculated within 100m of each station. This buffer was chosen pragmatically to approximate the detection radius of point counts and encompass a 3×3 pixel window, reducing the influence of anomalous pixel values compared to a single-pixel extraction. These 26 variables provide a high-dimensional description of local vegetation reflectance. To reduce redundancy, we used principal component analysis to condense the 26 variables, and the first five components, explaining 96% variation in the data, were used. PC1 (50% of variance) and PC2 (26% of variance) were not dominated by any single variable or group. The maximum contribution of any variable was 7% for PC1 and 11% for PC2, with most variables contributing more modestly. Variables with near zero loading in PC1 tended **to** contribute more to PC2 and vice versa. Thus, the components represent broad combinations of vegetation indices and reflectance bands. The distribution of the five principal components and the proportion of deforestation at three scales are plotted in **Error! Reference source not found**., and the relationship between the first two principal components with environmental descriptors is shown in **Error! Reference source not found**.

**Fig 2:**
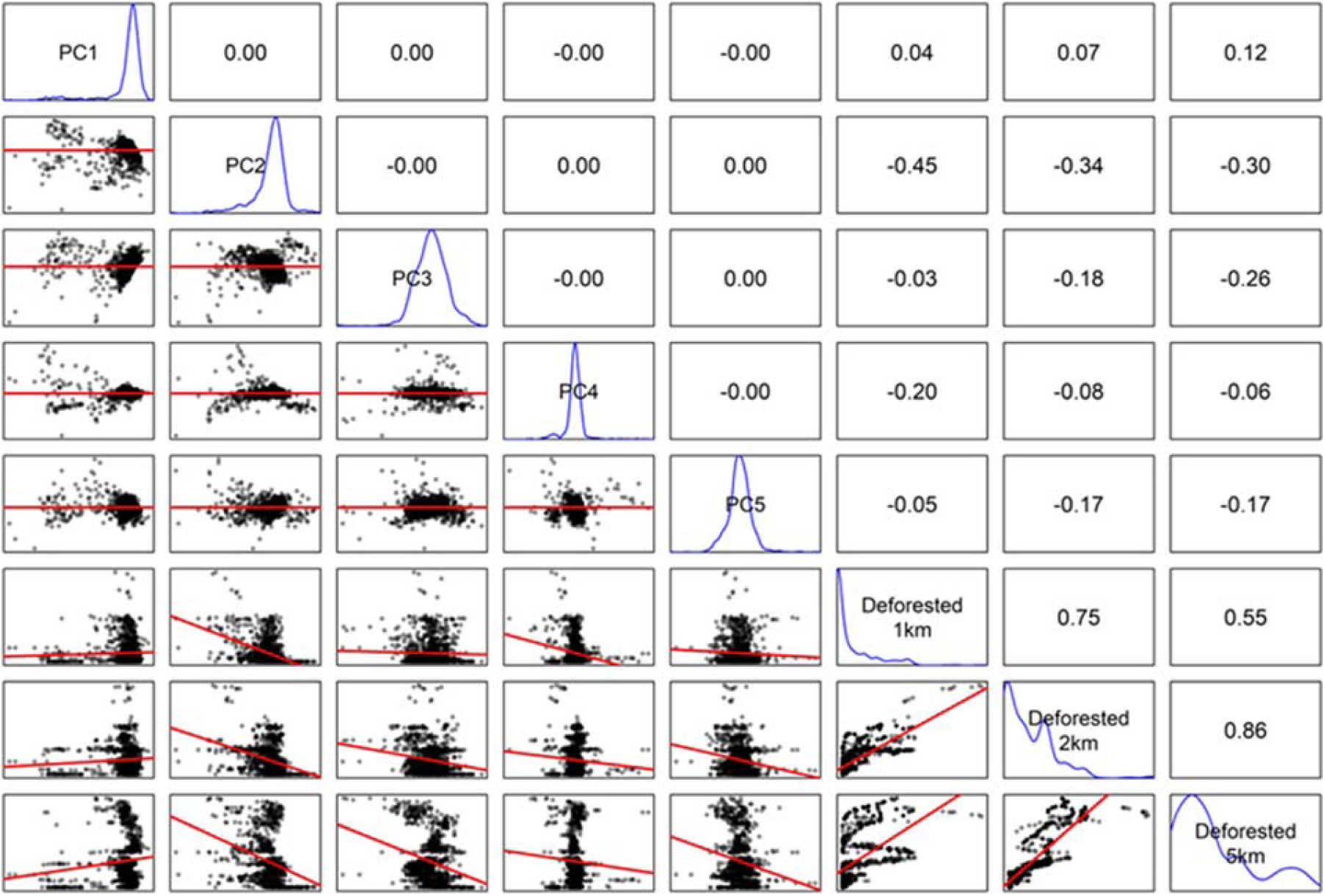
Pairwise relationships among predictor variables used in EO models. Lower panels are fitted with linear trend lines (red). Diagonal panels show variable names and their density distribution (blue). Upper panels show Pearson correlation coefficients.

**Fig 3:**
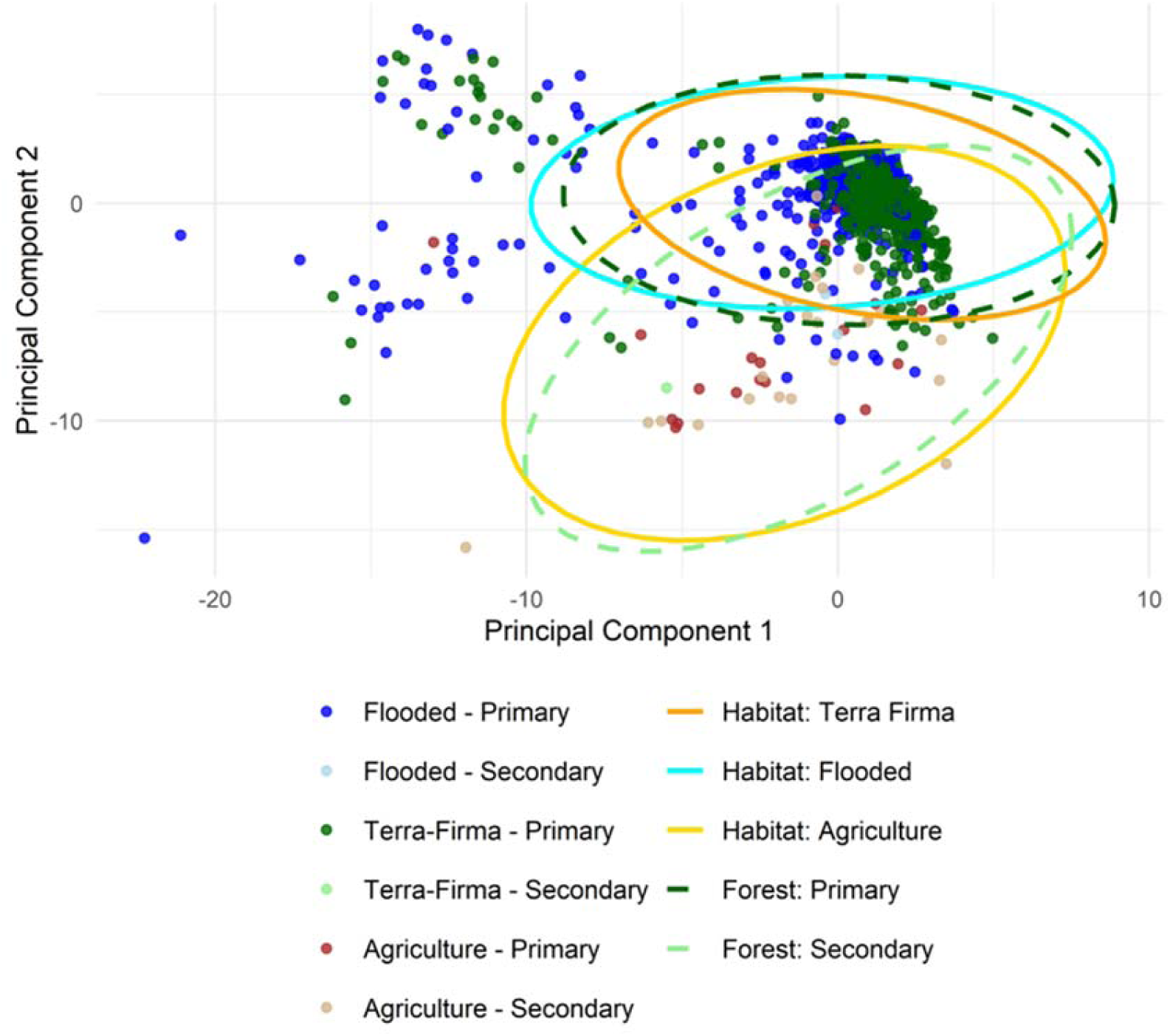
The first two principal components, representing 76% of the variation in the RS data, plotted against each other and showing their relationship with the measured habitat types. Each point represents an individual survey and is coloured based on the combination of habitat (floodplain forest, terra-firma forest, agriculture) and forest type (primary or secondary). Ellipses represent the 95% confidence interval around the mean for each categorical group.

### 3.4 Analysis

To quantify species detection and assess whether bird community composition can be predicted as a function of habitat or RS descriptors, we used spatial factor multi-species occupancy models. Observation depends on true presence (1 = present, 0 = absent; assumed constant over the year of surveillance), and the probability of detection. Detection can be influenced by survey method. Mist nets at ground level tend to under-sample canopy dwelling, large or less mobile species (49). Capture rates also vary with the type of net used (e.g. mesh size), and netting duration. Point counts detect vocal species well, but may under-sample silent or secretive species, and their effectiveness depends on background noise, weather, and observer skill (50,51). Perfect detection of all species is essentially impossible. Non-detection can bias covariate estimates and predictions of community composition (52). However, occupancy models account for imperfect detection by estimating species occurrence at each survey station, whether observed or not (53). When adequately specified, occupancy models can estimate detection probabilities per species and station, incorporating the influence of survey-specific covariates, such as survey method or time of day (54). After fitting the model and estimating species detection and occurrence probabilities, the methods of Guillera-Arroita and Lahoz-Monfort (2012), were used to calculate the power of the study. The term “power” is the probability of detecting a proportional reduction in occupancy between surveys conducted at different times. We then calculated the survey effort required to achieve a mean power across all species of 0.7, to detect a 50% occupancy reduction. Although 0.8 is commonly used, our low detection rates made 0.7 a more feasible target that falls within the test ranges (55). We also assessed how detection power of various occupancy reductions varied with the number of stations and replicate surveys, to guide survey regimes for future monitoring.

Four models were compared: one fitted with habitat variables, one with principal components from RS reflectance, one with no occurrence variables to act as an intercept-only baseline, and one RS model re-fitted without spatial factors to assess their influence. All models included survey method (point or net) and time of day (a.m. or p.m.) as covariates for the probability of detection. A summary of all detection and occupancy covariates, and the models in which they were included, is provided in Table 1. Although some stations were surveyed up to 21 times, 99% of all stations were surveyed 12 times or less. To improve the efficiency of model fitting and reduce the computational resources required to run them, we fitted the modes using a maximum of 12 replicates per station.

**Table 1:** Summary of covariates considered in the multi-species occupancy models.

| Covariate | Type | Description | Source / Processing | Models used in |
| --- | --- | --- | --- | --- |
| Land class | Categorical | Floodplain, terra firma, agricultural | MapBiomas Peru Project, Collection-1 (2013) (39) | Habitat model |
| Forest type | Categorical | Primary vs. secondary forest | Global Forest Change (2000–2019) (14) | Habitat model |
| Deforestation (%) | Continuous | Proportion of non-primary forest within 1, 2, and 5 km radii around stations | Global Forest Change (2000–2019) (14) | Habitat model |
| Reflectance indices | Continuous | NDVI, EVI, NBR, NDWI, TCB, TCG, TCW; means and SDs within 100 m | Landsat 5/7/8 annual median composites, Google Earth Engine | RS model → PCA |
| PCA components (PC1–PC5) | Continuous | First five PCs of 26 reflectance variables (96% variance explained) | Derived from Landsat indices above | RS model |
| Survey method | Categorical | Point count vs. mist-net | Field survey protocol (Fauna Forever) | All models (detection) |
| Time of day | Categorical | Morning vs. afternoon | Field survey protocol (Fauna Forever) | All models (detection) |

Detection covariates were included in all models. Occupancy covariates varied by model: habitat descriptors (land class, forest type, deforestation), RS reflectance (PCA components), or none (intercept-only).

Detection and occupancy covariates were modelled without interactions. Model outputs are on the logit scale, with baseline categories set alphabetically (a.m., net, agricultural, primary). Posterior estimates for other categories (e.g., p.m., point counts, secondary forest) were obtained by adding the relevant coefficient to the baseline and applying the inverse-logit transformation. We then summarised these posterior values to report the median and 95% credible intervals (2.5% and 97.5% quantiles). Reported detection and occupancy values in the Results are therefore directly derived from the posterior distributions.

Models were fitted using the Bayesian occupancy model package, “spOccupancy” (56). The “spOccupancy” package enables the modelling of all species simultaneously, and accounts for autocorrelation between species. We included three spatial latent factors to account for residual spatial autocorrelation in occupancy probabilities. The choice of three factors was a pragmatic one, guided by recommendations in the spOccupancy framework that a small number of factors (typically 2-5) provides sufficient flexibility to capture spatial structure while avoiding overfitting and excessive computational cost. Each factor provides a decay rate (phi), with the distance beyond which autocorrelation is negligible calculated as 3/phi. Models were fitted with 50,000 iterations, discarding the first 25,000 as burn-in, and thinning the rest by 50 to produce 500 posterior samples. Each posterior sample provides a probability of occurrence (psi) for each species at each station and a latent presence/absence (z = 1/0) calculated from this probability. Additionally, detection probability (p), given presence, is provided per species per station for a single survey. As spatial models often show poor multi-chain convergence, and given memory constraints, we chose to run a single-chain model. Convergence was assessed visually, with Geweke diagnostic (75% of parameters had z<1.96) and effective sample sizes (median ∼ 274 across all parameters), following spOccupancy guidance (57), all of which indicated adequate convergence. Models assumed no false presences and treated the entire study as a single closed season. Model goodness-of-fit was measured using posterior predictive checks to calculate a Bayesian p-value, based on the differences between chi-squared values of observed and predicted data per species, grouped across stations,(58,59) Models were also compared using the widely applicable information criterion (WAIC) (60) and metrics of predictive performance. Performance was evaluated at the species-level, using area under the receiver operating curve (AUC), and at the station community-level, using species accumulation and Bray-Curtis’s dissimilarity of community composition.

Although model performance differs between training and withheld data, evaluation on independent datasets is rarely undertaken for occupancy models (61). We did so here by withholding a validation dataset comprising two stations per field centre (24 centres in total), each surveyed at least three times, totalling 5% of the data. This ensured representation across all centres while retaining sufficient training data even in the smallest centres (minimum 4 stations). Validation stations were randomly chosen from those with at least 3 survey replicates, so their observed communities were well characterised for comparison with model predictions. To validate model performance, we incorporated both occurrence and detection probabilities when comparing predictions with observations. Each of the model’s posterior samples predicts the occupancy probability (psi) per species per site, and the latent occupancy (z = 1/0) representing presence or absence. Detection probability (p) per species and site in a single survey was predicted, and this remained constant across posterior samples. Total detection probability per species and station was calculated as 1 – (1 – p)^n^, where n is the number of replicate surveys performed. This detection probability generated a binomial detection/non-detection (1/0) event per species per station. Latent detection values were then multiplied by occurrence (psi) and latent occupancy (z) for each species and station per posterior sample. Thus, model performance was based on the predicted probability of detection rather than occurrence in the validation data.

## 4 Results

### 4.1 Model fit

Posterior predictive checks gave all models Bayesian p-values between 0.467-0.477, indicating a good fit to the data. The model with the lowest WAIC value, thus indicating it was the best-performing model, was fitted with RS reflectance data and included spatial factors, followed by the covariate-free intercept model, the RS reflectance model fitted without spatial factors, and the highest WAIC indicating the worst performance was from the model fitted with environmental descriptors.

The probability of detection (_p_) was consistent across all models. Surveys conducted in the morning using net counts had a posterior median species-level _p_ = 0.022, 95% CI [0.001, 0.179]. When conducting surveys in the afternoon, median detection probability significantly lower at _p_ = 0.015, 95% CI [0.001, 0.018]. When using point rather than net surveys, the median _p_ increased significantly to 0.065, 95% CI [0.000, 0.389]. These results indicate that detectability depends strongly on method and time of day, with point counts and morning surveys yielding higher overall detection rates. However, point surveys did not significantly increase detection for three of the nine Orders represented. Those three were Caprimulgiformes (nightjars), and Coraciiformes (kingfishers and hornbills), and Galliformes (ground dwelling birds).

The posterior mean probability of occurrence across all species in primary forest and an agricultural habitat, was 0.291, 95% CI [0.211, 0.375]. In secondary forest, the mean occupancy was higher than in primary forest, rising to 0.47, 95% CI [0.36, 0.61]. In comparison to agricultural landscapes, mean occupancy in floodplain increased to 0.36, 95% CI [0.30, 0.47], and in terra-firma mean occupancy increased to 0.37, 95% CI [0.30, 0.47]. This indicates that secondary forests supported more species on average than primary forests, while agricultural habitats supported fewer. Mean occupancy was not shown to vary significantly with deforestation within a 1-2 km radius, but increased by 0.057, 95% CI [0.008, 0.108] with each 10% increase in deforestation within a 5km radius. This suggests that bird occurrence responded more strongly to landscape-scale forest change (5km) than local variation (1-2km).

When using RS data, the model suggested that spatial autocorrelation extended to 5km, 95% CI [3.3km, 7.5km] for one factor and 7.5km, 95% CI [4.3km, 15km] for each of the other two factors. When using habitat data, the model suggested that spatial autocorrelation extended to 2km, 95% CI [2km, 2.3km] for one factor, to 30km, 95% CI [3.75km, ∞] for another, and the third factor was not significant. These factors represent unmeasured variables that influence the occurrence of species, and fade with distance. Although we do not know what is driving this spatial structure, it can be influenced by anything from local vegetation cover to regional climate or land use. Significant spatial autocorrelation confirm that patterns exist beyond those explained by the known environmental variables.

### 4.2 Model validation

The predictive performances of models followed the implications made by their WAIC values, and the model that produced the best predictive results was also fitted with RS reflectance data and included spatial factors. When assessed on training data, this model had a mean AUC of 0.88 (SD=0.06) and when predicting to independent validation sites, had a mean AUC of 0.68 (SD=0.18; range=0.16-0.99). Species for which predictive performance was considered good (AUC>0.7) (62) are hereafter referred to as high-AUC species, and this model had 49 high-AUC species, the highest of any model. The performance of all four models is shown in Table 2**Error! Reference source not found.**. Species taxonomic order was not shown to have any correlation with its AUC.

**Table 2:** The fit and predictive performance of the four models analysed.

| Model Covariates | Spatial Factor Included | WAIC value | Delta WAIC | Mean Explanatory AUC | Mean Predictive AUC | Number of high-AUC species |
| --- | --- | --- | --- | --- | --- | --- |
| RS Reflectance | yes | 66773 | 0 | 0.88 | 0.68 | 49 |
| RS Reflectance | no | 68620 | 1847 | 0.87 | 0.62 | 31 |
| Habitat Descriptors | yes | 69432 | 2659 | 0.87 | 0.58 | 21 |
| None | yes | 66995 | 222 | 0.8 | 0.66 | 43 |

Low WAIC values and high AUC values represent better performances. It can be seen that the best performing model in all categories is fitted with RS reflectance data and includes spatial factors.

The predicted species richness of survey stations (6.86 ± 5.1) was close to the observed average richness (6.00 ± 3.6), but when extended to the entire training dataset the model predictions suggested that it would take much longer to detect the total number of species included (**Error! Reference source not found.**(A)). The mean predicted richness of test stations (9.94 ± 3.8) was also comparable to the observed mean richness (10.82 ± 5.3) and indicated a similar accumulation of species, both across stations (**Error! Reference source not found.**(B)), and with increasing replication within a station (**Error! Reference source not found.**(C)). The species accumulation curve in **Error! Reference source not found.**(C) come from one station that was surveyed 12 times and illustrates the extent to which observed accumulated richness greatly underestimates the total richness expected to be present at each location. A consequence of this was that only a fraction of the species that occur at a station were observed, and therefore the average Bray-Curtis dissimilarity between the list of birds observed and those predicted was 0.8 (± 0.1). In locations with higher observed richness, the surveyed total represented a greater proportion of the true expected total, and observed species richness was strongly negatively associated with predicted-observed community dissimilarity (R^2^=0.68, p<0.001) (**Error! Reference source not found.**).

**Fig 4:**
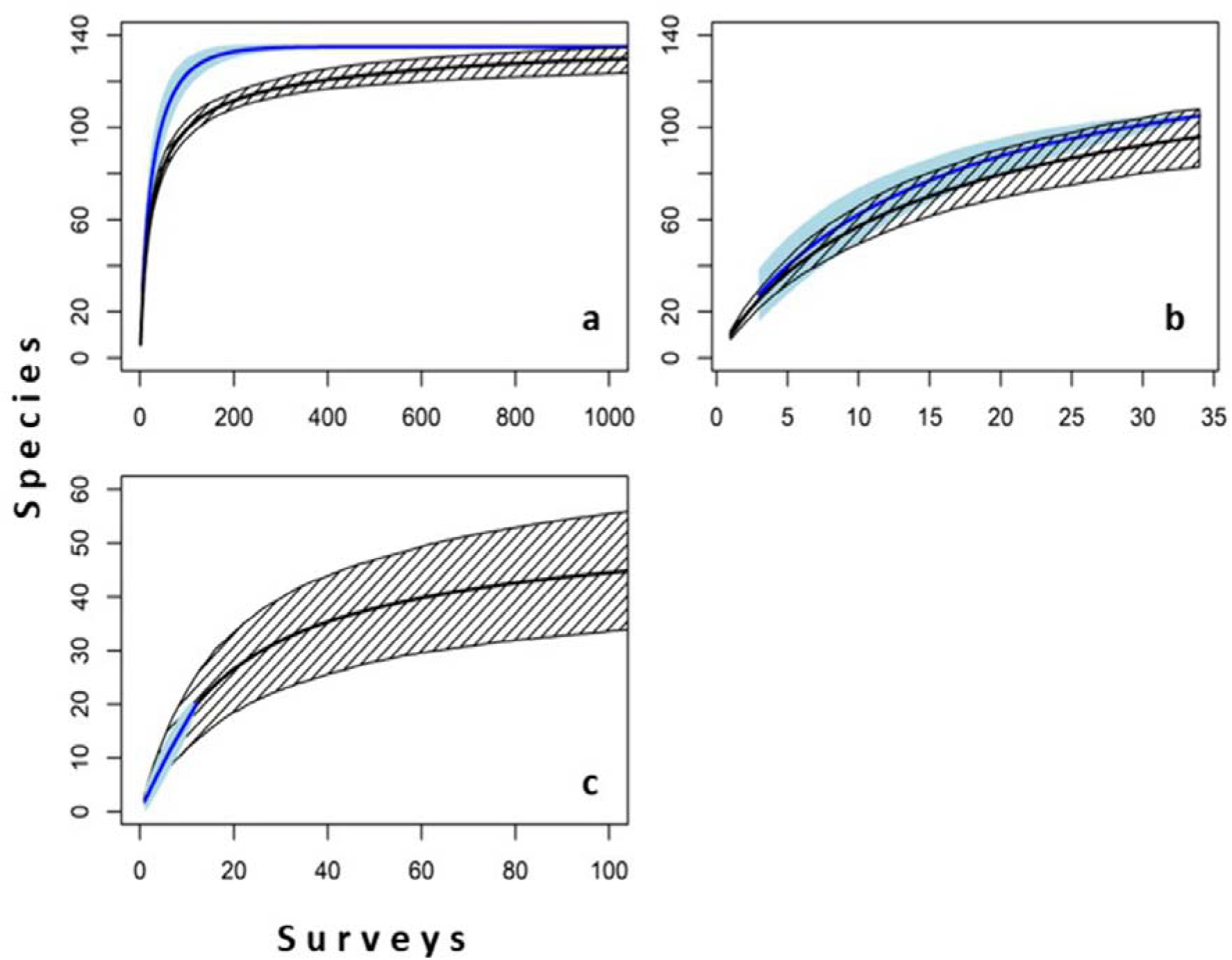
Observed (blue) and modelled (black) species accumulation (±2SD) with increasing numbers of surveyed stations, (a) within the training data, (b) set aside for validation, and (c) based on an increasing number of replicate surveys conducted at a single survey station.

**Fig 5:**
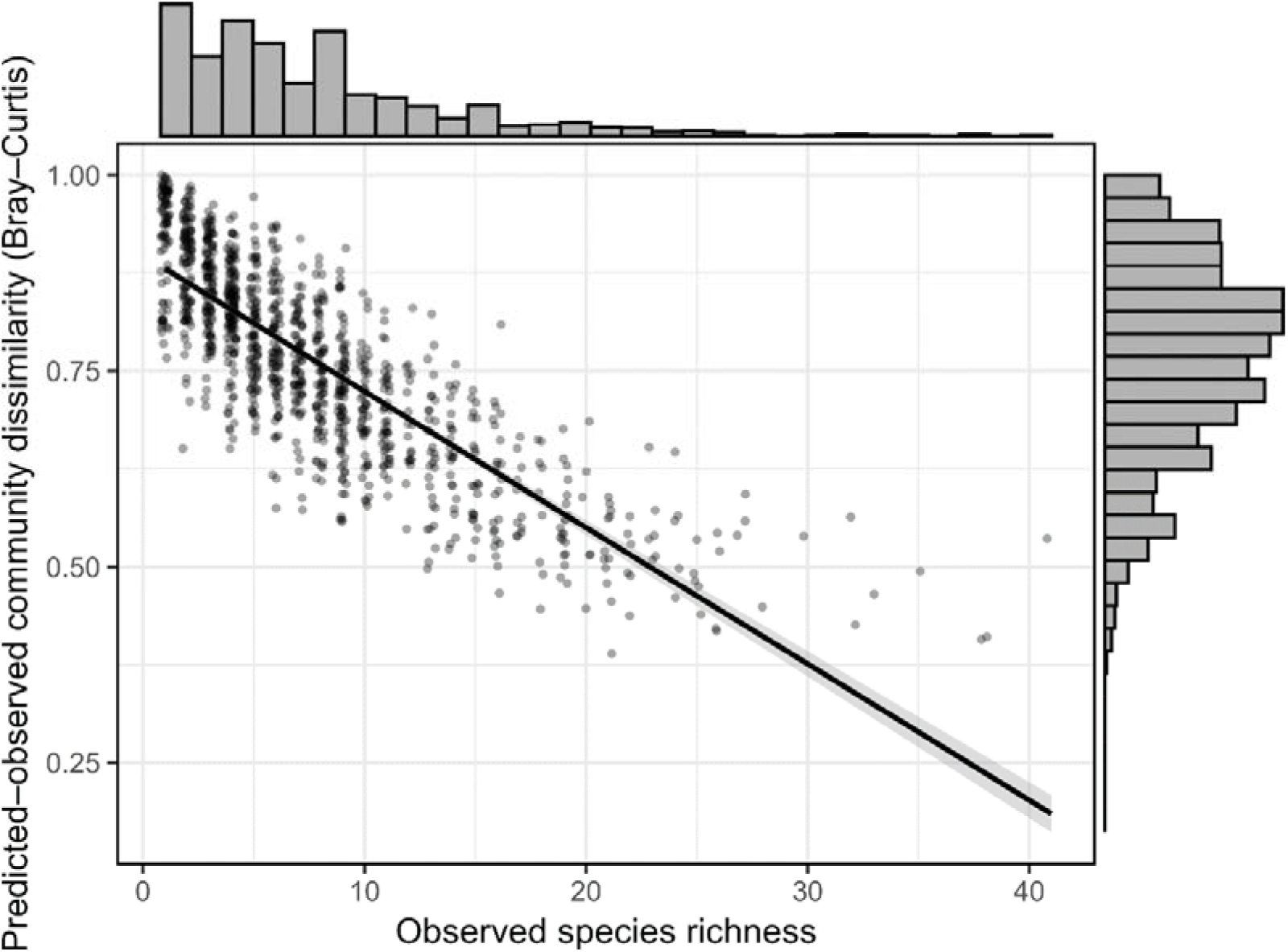
Relationship between observed species richness and dissimilarity between predicted and observed community composition (Bray-Curtis) across all sites. A fitted linear regression line (black) with 95% confidence interval (dark grey band) is shown (R =0.68, p<0.001). Marginal histograms show the distribution of each variable.

### 4.3 Power to detect change

As a result of low detection probabilities and low mean replicate surveys, it was ascertained that there is a mean probability across all species of 0.20 (±0.24 SD) of detecting a 50% reduction in occupancy in a subsequent survey. There is a 70% or greater probability of detecting a 50% reduction in only five of the 135 species modelled. Thus, if the desire is to determine where occupancies are changing by surveying the same area in the future, the current data cannot provide robust evidence.

illustrates how different numbers of survey stations and replicates per station affect the median power to observe a reduction in bird occupancy. Each panel represents a different proportional change that is sought to be detected, and each y-axis represents the probability of detecting that change in at least half the bird species. The x-axis is the total number of individual surveys that a study is able to undertake (i.e. one replicate at one station is one survey). Each line represents a different number of sites that the total number of surveys are divided among. For every probability of detecting the panel’s proportional change (y-axis), the farthest left line represents the fewest total surveys required, and the most efficient number of sites to survey. For example, if our aims were to have a 70% confidence (0.7 on the y-axis) that we could detect when the occupancy of at least half the bird species had declined by 50% (panel C), the chart suggests that the fewest surveys required would be ∼10,000 and spread between 100 and 500 sites. Calculations suggest that the requirements would be three replicate surveys at nearly 50,000 stations, or, if all stations were surveyed 27 times, only 447 stations would be required.

**Fig 6:**
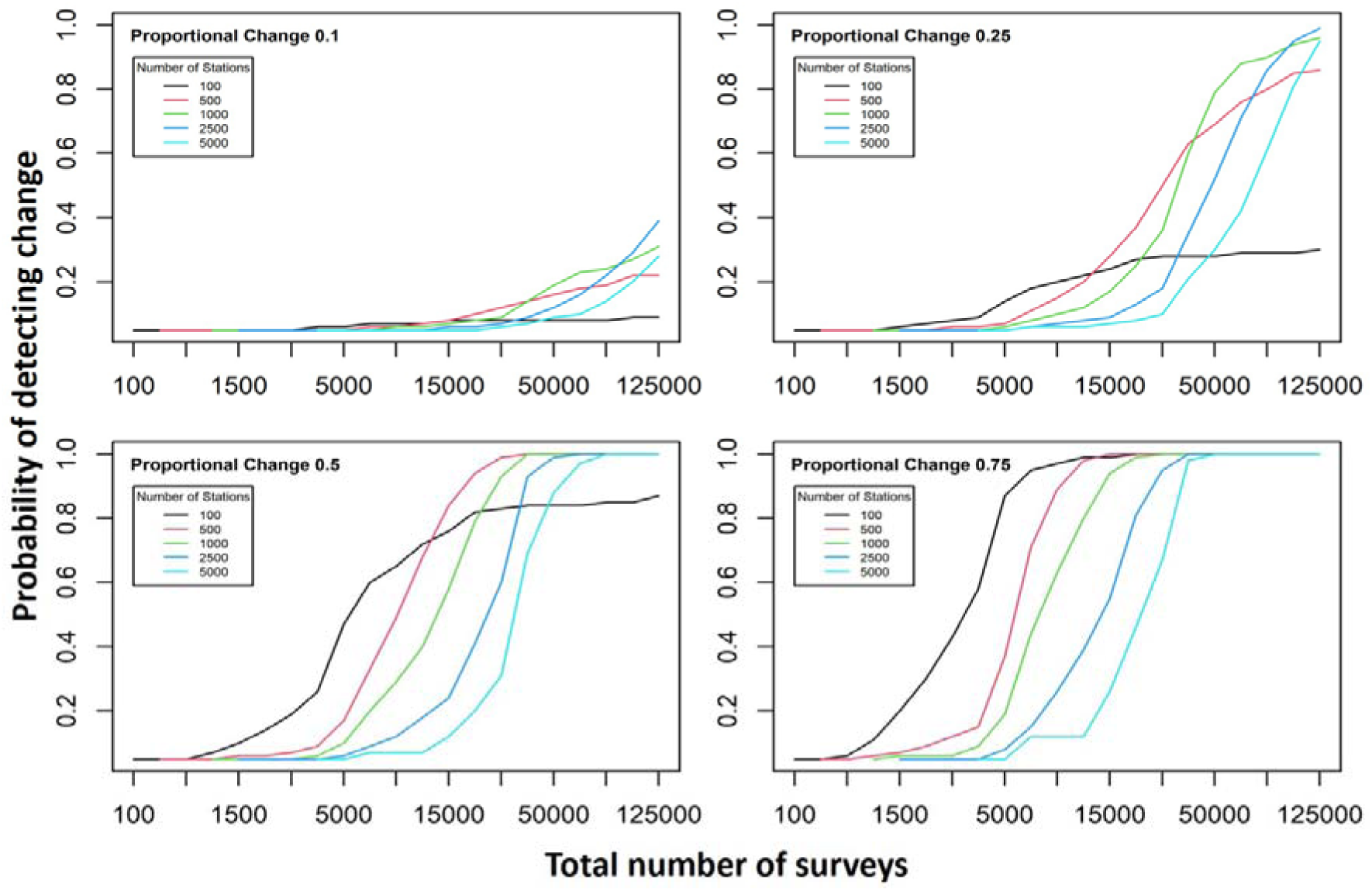
The influence of survey effort on the probability of detecting a reduction in occupancy in future surveys. Survey effort was calculated as the number of stations surveyed * number of replicate surveys per station. Each line represents a different number of stations and was calculated as the mean value across all 135 species modelled. Each chart represents a different proportional reduction in occupancy.

## 5 Discussion

Using over 3,000 bird surveys conducted in south-eastern Peru, we found that remotely sensed satellite data performed slightly better than categorical landcover descriptors in predicting bird community composition. This advantage was modest and specific to our study region, and further testing would be required before generalising to other tropical systems. We also highlight the low detectability of many species, which complicates biodiversity monitoring in rainforests. Although the survey design provided valuable insights into bird communities, it was not optimised to detect temporal change. Our results can guide more efficient survey strategies, leading to enhanced species detection, better capturing community composition, and strengthening the performance of RS models in predicting bird communities across landscapes.

### 5.1 Land class and surface reflectance

Studies differ on how forest and farmland affect bird species richness (63,64). We found more species in both floodplain and terra firma forests than in agricultural land, partially aligning with evidence that species richness can increase at forest boundaries, particularly where edges are sharp, such as where forest and agricultural land meet (65,66). Furthermore, the greater stratification and canopy complexity often found in primary forests typically support more specialist, range restricted, and endemic bird species compared to secondary forests (67–69). However, our analysis suggested that secondary forest had a greater species richness, consistent with findings that generalist and disturbance tolerant species as well as species that thrive in more open canopy environments are drawn to those areas, especially when they are connected to primary forest, thereby increasing species richness (70). Greater visibility, vocalisation and bird density in open secondary forests may also lead to more detections (71–73). The low mean number of replicates in our study could make secondary forests appear richer in species than primary forests, due to improved visibility rather than true richness. Secondary forests in Amazonia typically occur as mosaics of regenerating lands, agricultural edges, and disturbed forest patches. These mosaics can provide food resources and structural heterogeneity that are especially favourable to generalist Amazonian bird species (74). However, our data cannot disentangle these ecological drivers from detectability effects, and both are likely contributing to the higher richness observed.

Secondary forests require the initial loss of primary forests, and localised deforestation can increase bird species richness. Moderate primary deforestation can increase richness of neighbouring forest as displaced individuals move into it (75). It can also increase habitat heterogeneity, promoting colonisation by generalists and disturbance-tolerant species in the deforested area (75). Our findings that deforestation within a 5km radius increased mean occupancy support this. At this scale species dispersal, habitat connectivity, and edge effects shape community composition (76,77). In contrast, no effects were found at smaller distances, suggesting insufficient heterogeneity at that range. We defined deforestation as loss of primary forest, but deforested areas may have regrown into secondary forest, which also showed high species richness. Misclassification could arise from pre-2000 deforestation or from the omission of water bodies from analysis. Within floodplain forests, water levels can fluctuate seasonally (78) and edge effects may occur at interfaces between large water bodies and forests (79). These classification limitations should be considered when interpreting differences in species occupancy across forest types. bird detectability may have varied due to seasonal fluctuations, which are most pronounced in forest fragments and secondary forest (67,80,81). Surveys were performed by multiple observers with varying levels of experience, which can significantly influence bird species detectability (82). Consequently, some variation in detection probabilities may be attributed to differences in surveyor rather than actual ecological differences.

Surface reflectance diversity has been used to predict tropical forest and bird diversity (83) and bird species richness found within them (84) and our findings support this. RS data performed as well as, if not better than, land class in predicting bird communities (85,86). Image seasonality and resolution matter for modelling bird communities in temperate regions (85,86). However, persistent cloud cover in our study area required annual composites, and mean reflectance values calculated over thousands of square metres make finer resolution imagery unlikely to improve results.

Despite these challenges, our study has shown that RS data have the capacity to improve on land class in predicting bird communities in tropical forests. Two broad land-class groups could be differentiated by plotting the first two PCAs: secondary forest/agriculture and primary forest/terra-firma/floodplain. Within those two groups, secondary forest, defined as land deforested since 2000, had a large overlap with agricultural land, suggesting that deforested land here tends to become agricultural rather than regenerating into secondary forest (87,88). In the other group, primary forest overlapped more with terra-firma and floodplain than with agriculture, though sizeable overlaps between all groups existed. This may be explained by discrepancies brought about by course scale, out of date information or human error in habitat classification (89–91). Additionally, land classes may oversimplify or fail to capture the finer scale variation seen by RS data. Localised secondary descriptors of structural variation, such as deadwood volume and tree girth, can also influence reflectance values, and when included in conjunction with broad habitat type, have been shown to define species richness far better than habitat type alone (92). This shows that RS data better capture gradual changes in vegetation structure than can predefined classes. The challenge for future research will be in deriving RS variables that can describe more ecologically meaningful variation and differentiate between less obvious variations in surface structure.

### 5.2 Realities and practicalities of observing change

Whether using habitat or RS variables, model success depends on the quality and quantity of the data with which they are trained (93,94). Low detection probabilities suggest substantial effort is needed to detect most species, raising concerns about data sufficiency. However, of the approximately 760 bird species thought to occur in the Tambopata area (95), our study recorded nearly half. This, in terms of both number and proportion of species identified, aligns with findings from other surveys in tropical forests (96–98). Low detection rates likely caused individual surveys to underestimate community composition, making landscape-wide communities appear more distinct than limited survey observations suggest. Nevertheless, the species modelled exhibited patterns of grouping based on both forest habitat descriptors and RS variables, suggesting that they represented more than just random observations. This indicates that, despite describing approximately a half of the full community, our results were reliable.

Increased sampling effort revealed more of the local community and improved similarity between predicted and observed communities. Thus, improving detection rates is essential to increasing the proportion of the community observed and improving the predictive output of models. Increasing cumulative detection rates to 80% has been shown to negate the importance of detection on model performance (99). Yet we have shown that it would require a huge effort to reliably observe changes in occupancy for many species. Thus, efficient and accurate surveys are critical to provide high quality data on which to train RS models, especially as ongoing surveys are needed to monitor the effects of conservation efforts on habitat change over space or time. The type of survey conducted affects detection rate, and we agree with Mulvaney & Cherry, (2020) that point counts increase mean detection over net counts. However, these findings are not universal, as it has been shown that within tropical forests, the most effective survey method varies with ecosystem, such as whether the study is within lowland or cloud forest (97). Similarly, transect counts and canopy counts add further depth to the proportions of small-bodied species, canopy and mid-storey species that are detected (82,98). Due to the relatively low community overlap observed between methods found in our study, and aligning with other studies, we would suggest that the use of multiple methods is necessary to improve overall diversity assessment (82,97,98). Further to the survey methodology lies the challenge of recruiting appropriately trained surveyors who are familiar with the bird species of the area, which is a necessity in conducting effective surveys in complex and rich ecosystems such as tropical forests (82). Our results suggest that to effectively detect changes in community composition, it is necessary to perform far more replicate surveys than were performed (82).

### 5.3 Future potential

Forest degradation in Peru has been shown to increase the amount of carbon released into the atmosphere by almost 50% (101). Protecting forests and carbon sinks has understandably been central of tropical conservation, and government policies and voluntary carbon offset conservation efforts such as REDD+ have successfully slowed the rate of deforestation in recent years and have played a role in mitigating climate change (102). To ensure that the implementation of carbon conservation efforts, such as REDD+, also have a positive impact on biodiversity, ecological monitoring of forests should be aligned with the conservation objectives of the area (103). While REDD+ has had some success in protecting forests globally, the level of success in the Amazon may be overstated due to the overestimation of baseline deforestation (104), highlighting the need to validate the costs of voluntary carbon offset credits. Degradation impacts are not fixed and can be improved by restoration or worsened by repetition and should be measured at a landscape rather than patch scale (105). Measuring community diversity at landscape scales may require impractical levels of monitoring, but patch scale changes in habitat should in principle be observable by Earth observation. Should sufficient, consistent, reliably representative biodiversity data be available to calibrate models fitted with RS data, biodiversity responses to changing land use can be identified. Policy and management should judge the results of practices at the point of implementation rather than judge the practices conceptually as a whole, and by mapping species diversity in conjunction with other measures of landscape health, trends in forest degradation and regeneration over time can be identified (105).

Despite the challenges posed by generally low detection rates, our study demonstrated that the occupancy of a third of the bird species with sufficient detections in the Peruvian Amazon can be predicted with reasonable confidence. However, low detection rates limit model capacity to detect community change over time. While RS models may not yet be sufficiently robust for ongoing monitoring, reporting and verification, there is no evidence that field-based methods alone will be able to do this at the scale required. While surveys can identify spatial variations, detecting temporal change requires detailed count data at a level that may be impractical (106). Our power analysis revealed that with the current monitoring structure, even if a species population were to half, we would fail to detect this four out of five times. The suggestion that declines of many species may be substantial before being reliably detected has major implications for conservation monitoring using existing survey design. Improving detection rates will increase the power of future surveys to detect change. Given that resources are limited, future surveys should focus on fewer locations, prioritised across an ecological gradient of interest, with a higher number of replicates; results suggest at least 20 per site. This is particularly critical as even basic inventories of avian diversity across much of the tropics are lacking (82). Establishing baseline datasets is essential for enabling future repeat surveys, which could provide valuable insights into how and where tropical bird communities are changing (107). Building on the findings of this study by refining survey design and aligning research with specific conservation objectives will be key to improving biodiversity monitoring in the region so that decisions are based on detectable trends rather than snapshots of biodiversity (108). Incremental improvements in temporal monitoring are most likely to come from combining these refinements in survey design with successive RS image series. By aligning more robust field baselines with RS predictors, it will become possible to track biodiversity change gradually across space and time, even in data-limited tropical landscapes.

## Declarations

### Ethics Statement

The bird occurrence data used in this study were collected independently by co-authors C.A.K. and C.K. as part of long-term monitoring within the Tambopata National Reserve, Peru. Data collection was conducted ethically and in accordance with local research regulations.

### Authors’ Contributions

Author contributions: A.C.S., A.B. and I.R.H. conceptualised the study; C.A.K and C.K. performed independent field surveys and provided bird sighting data; A.C.S. sourced and calculated all remotely sensed variables, performed all modelling and analysed results; A.C.S. wrote the manuscript with reviews and edits suggested by A.B, I.R.H.

